# Tau hyperphosphorylation on T217 in cerebrospinal fluid is specifically associated to amyloid-β pathology

**DOI:** 10.1101/226977

**Authors:** Nicolas R. Barthélemy, Randall J. Bateman, Philippe Marin, François Becher, Chihiro Sato, Sylvain Lehmann, Audrey Gabelle

## Abstract

**Introduction:** Modification of CSF tau phosphorylation in AD remains controversial since total-tau and phospho-tau levels measured by immunoassays are highly correlated.

**Methods:** Stoichiometry of phosphorylation of CSF tau was monitored on five sites by mass spectrometry. We compared 50 participants with AD at mild to moderate stages, others tauopathies and controls then confirmed our results in a cohort of 86 participants cognitively normal or with mild cognitive impairment and stratified according to amyloid-β status.

**Results:** Changes in tau phosphorylation rates were observed in AD participants but not in other tauopathies. T181 appeared slightly hyperphosphorylated in AD. In comparison, T217 phosphorylation, was considerably modified. We demonstrated T217 hyperphosphorylation occurred systematically in amyloid positive participants even at preclinical stage (AUC 0.999). T217 phosphorylation change specificity overpasses other phosphorylated tau sites investigated in this study, but also CSF total-tau and p-tau levels.

**Discussion:** CSF T217 hyperphosphorylation defines a specific tauopathy status concomitant with ß-amyloidosis.

## 1. Introduction

Alzheimer disease (AD) is the leading cause of dementia worldwide (1). Its diagnosis and treatment remain extremely challenging in absence of specific and early biomarkers. Recent developments of cerebrospinal fluid (CSF) biomarkers and brain Positron-Emission Tomography (PET) imaging provided valuable tools to detect the pathognomonic AD brain pathologies of amyloid-β plaques and neurofibrillary hyperphosphorylated tau tangles. Currently, the CSF profile of AD patients is characterized by decreased amyloid-Beta 42 (Aβ42) and increased total and phosphorylated tau (p-tau 181) (2-4) measured with standard immuno-assays. This profile allows the discrimination of AD versus non-AD pathologies (5-8) and the detection of AD process many years prior to cognitive symptoms or complaint (9, 10). However, changes in CSF tau and p-tau are not specific of AD. Although the increased p-tau181 has been interpreted as a consequence of hyperphosphorylation due to tangle formation (11), CSF p-tau increases concomitantly with total tau. Brain studies indicate tau phosphorylation on numerous sites, but the diagnostic relevance of these additional phosphorylation sites in CSF has not been fully addressed. In particular, changes in tau phosphorylation at different sites related to specific neuropathological diagnosis and tau aggregates have not been compared in AD and other tauopathies such as Progressive Supranuclear Palsy (PSP), Cortico-Basal Degeneration (CBD), frontotemporal lobar degeneration (FTLD), other non-tauopathy cognitive impairment diseases or cognitively normal controls. Mass spectrometry (MS)-based methods are more relevant than immunoassays to assess changes in phosphorylation levels of specific sites independently of total tau levels as they allow independent quantification of phosphorylated peptides and their corresponding unmodified counterparts. Accordingly, correlations between phosphopeptide and unphosphorylated peptide slopes can be compared to evaluate the rate of phosphorylation. To quantify phosphorylated tau isoforms in CSF (12) we used an innovative targeted high-resolution MS (HRMS) method targeting tau phosphorylated peptides in the mid-domain of the protein sequence, which is the most abundant domain in CSF (13) and is phosphorylated on numerous sites in brain tau and AD tau aggregates (14).

We analyzed tau phosphorylation on T181, S199, S202, T205 and T217 in CSF using a cohort comprising probable AD (with high level of significance according to the NIA criteria) with mild to moderate stage of the disease, other neurological disorders including tauopathies and cognitively normal controls. Then, we validated our results in a second cohort comprising cognitively normal individuals and patients with mild cognitive impairment stratified by their amyloid status based on CSF Aß42/40 ratio and PET-PIB imaging. This validation allowed us to highlight the potential of CSF pT217 for AD diagnosis and to establish correlations between pT217 and amyloidosis underlying tau modification at an early stage of the disease.

## 2. Methods

### Discovery step

#### 2.1 Participants (Discovery)

We selected CSF samples stored at the Montpellier CSF-Neurobank (#DC-2008-417 of the certified NFS 96-900 CHU resource center BB-0033-00031, <u>www.biobanques.eu</u>) (SI Appendix, Table S1). CSF Aβ42, Aβ40, total tau and p-tau181 levels were measured using the standardized commercially available Innotest^®^ sandwich ELISA (Fujirebio) (32). Fifty participants followed at the Montpellier Memory Research and Resources Center were analyzed. For the diagnosis validation, a two step-procedure was performed as described in Supplemental Text S1-S2. Thus 10 patients with probable AD with high level of evidence of AD pathological process (mean age = 75.8 ± 9.9; sex ratio 8/2 F/M) according to the NIA diagnosis criteria (15) were compared to 40 non-AD disease (mean age = 69.0 ± 12.2; sex ratio 13/27 F/M) including 5 FTLD, 9 LBD, 7 PSP, 1 CBD, 7 adult chronic idiopathic hydrocephalus (ACIH), 2 possible AD etiologically mixed presentation or AD with concomitant cerebrovascular diseases (AD with CVD), 2 vascular dementia (VD), 1 with brain metastasis (BM) and 6 cognitively healthy controls). LC-MS analyses were performed blinded to clinical diagnosis.

#### 2.2 CSF tau chemical extraction (Discovery)

CSF samples (450 μl) spiked with 15N-tau-441 (final concentration 100 fmol/ml) were extracted as described previously (12). After precipitation with perchloric acid, acidic supernatant was extracted by solid phase extraction, dried then digested with trypsin. The digest was acidified with 5 μl of 10% acid formic. For phosphopeptides quantitation, AQUA phosphopeptides labeled at the peptides C-terminal residue TPP APKp TPP S S GEPPK (pT181), SGYSpSPGSPGTPGSR (pS199), SGYSSPGpSPGTPGSR (pS202), SGYSSPGSPGpTPGSR (pT205) and TPSLPpTPPTREPK (pT217) (Thermo Fisher Scientific, Ulm, Germany) were spiked to achieve individual concentration of 100 fmol/ml per labeled phosphopeptide in the final extract. Extracts were analyzed by LC-MS/HRMS as described in Supplemental Text S3. Quantitation, extraction of MS/HRMS transitions and integration of extracted ion chromatograms were performed using Xcalibur 2.2 (Thermo Scientific).

#### 2.3 Tau peptides and phosphorylated peptides quantitation (Discovery)

Stable isotope dilution (SID) method was used for MS quantitation by PRM. PRM transitions of phosphorylated peptides were quantified by comparison with labeled AQUA phosphorylated peptides (41). Heavy peptides released from 15N recombinant tau protein spiked prior the sample extraction, were used for unphosphorylated tau peptides quantitation as previously reported (12). Unmodified peptides were quantified using ratio with corresponding 15N-labeled peptides, as described previously (12). Measurement reproducibility was assessed using quality controls (QC) constituting of three CSF pools (t-tau concentration, assessed by ELISA: 144, 317 and 639 pg/mL respectively) both extracted four times independently and analyzed simultaneously within the cohort (Fig. 2 A-G). The response linearity of PRM transitions used for phosphorylated peptide quantitation was confirmed by reverse curves of phosphorylated AQUA peptides performed from pools of CSF extracts (Fig. S4).

#### 2.4 Participants (Validation)

Eighty-six participants cognitively normal (CDR=0) or with mild cognitive impairment were recruited from the Washington University in Saint Louis ADRC study previously reported by Patterson et al. 2015 with CSF and amyloid PiB-PET data available (26). Demographics of participants are described in Supplemental Table S5. This cohort included 29 amyloid positive and 47 amyloid negative participants according to the results of PiB-PET (considered positive above 0.18 used as cut-off) and CSF Aβ42/Aβ40 ratio measured by MS (considered pathological below 0.12 used as cut-off) and 10 cases with conflicting results between PET-PIB and CSF profiles (5 with a PiB-PET(+)/CSF(-) and 5 with PiB-PET(-)/CSF(+) amyloidosis profile). Seven CSF samples were extracted in duplicate and one in triplicate to assess variability.

#### 2.5 CSF tau purification using immunoprecipitation (Validation)

800 μl of CSF supernatant obtained after Aβ immuno-precipitation and storage at -80C was used for tau analysis. Thawed supernatants were spiked with 15N tau internal standard (5 ng per sample) and extracted using Tau1 immunoprecipitation. 5mM Guanidine, 1% NP-40 and protease inhibitor cocktail were added to the sample, then samples were mixed 3 hours at room temperature with 20 μl of sepharose beads cross-linked to Tau1 antibody. Beads were precipitated then rinsed with 0.5 M Guanidine and TEABC 25mM. Samples were digested with 400 ng of trypsin. AQUA peptides (Life Technologies, Carlsbad, California) were spiked to achieve individual quantity of 10 fmol per labeled phosphopeptide and 100 fmol per labeled peptide per sample. AQUA TPSLPpTPPTR (pT217) substituted the missed cleavage version used in the discovery cohort. Tryptic peptides were loaded on TopTip C18 tips, washed with 0.1% FA solution and eluted with 60%ACN 0.1%FA solution. Eluates were dried on speedvac. Samples were stored at -80C. Before LC-MS analysis, samples were resuspended in 25 μl 2%ACN 0.1% FA. Extracts were analyzed by nanoLC-MS/HRMS as described in Supplemental Text S4.

#### 2.6 Tau peptides and phosphorylated peptide quantitation (Validation)

Stable Isotope dilution MS quantitation using 15N labeled peptides was used to calculate absolute levels of unmodified peptides. Phosphorylation rate for each site was calculated by single point calibration comparing area ratio obtained on AQUA unphosphorylated and phosphorylated peptides counterpart to the area ratio measured for corresponding endogenous peptides. Phosphorylated peptide level was calculated by combining unmodified peptide level counterpart and rate of phosphorylation of the corresponding site.

#### 2.7 Statistics

Statistical analyses, including comparison of the slopes of regression lines, were performed using the GraphPad Prism software (7.0). Statistical significances between values obtained across investigated groups were calculated using non-parametric Mann-Whitney tests. Non-parametric Spearman’s rho rank correlation coefficients were used to assess the correlations between two series of values. Statistical significance was defined by p<0.05.

## 3. Results

### 3.1 Quantification of CSF tau phosphorylated peptides

To detect a set of very low abundance phosphorylated peptides from the tau mid-domain in CSF samples, we implemented a hypothesis-driven targeted-HRMS method (SI Appendix, Text 1). For the first time, this method allowed us to confidently detect tau phosphorylated peptides on residues corresponding to T181, S199, S202, T205 and T217 in the entire protein sequence (Fig. 1). To confirm phosphorylated site identification, we compared ion signals of the native peptides with synthetic isotope-labeled peptides spiked in CSF samples (Materials and Methods). Then, we quantified phosphorylated and unphosphorylated tau peptides by stable isotope dilution in CSF samples using from a cohort of 50 participants including 10 with probable AD with a high level of evidence of AD pathophysiological process (15). The description of all the participants is shown in Materials and Methods and SI Appendix, Table S1. In all CSF samples, peptides phosphorylated on T181 (pT181), S199 (pS199), S202 (pS202) and T217 (pT217) were detected. Phosphorylation on T205 (pT205) was mainly observed in AD samples (Fig. 2). pT181 exhibited higher concentration (mean 31.1 ± 22.3 fmol/ml, n = 49 in the cohort excluding BM), compared with peptides encompassing pT217, pS202, pS199 and pT205 (3.2 ± 4.9, 6.2 ± 3.6, 3.7 ± 3.0 and 1.3 ± 3.2 fmol/ml, respectively). Quantification of unmodified tau peptides and pT181-containing peptide by MS showed a strong correlation with t-tau and p-tau(181) levels determined by ELISA, respectively (SI Appendix, Fig. S1) (13). Three groups were identified based on tau, p-tau, phosphorylated/unphosphorylated peptide ratios measured by MS (SI Appendix, Fig. S2-S3): 1) the 10 probable AD patients, who show higher levels of all unmodified and phosphorylated tau peptides, 2) the 39 non-AD and cognitively normal controls with low or intermediate tau and p-tau levels, and 3) one outlier that corresponds to a patient with the Brain Metastasis. To understand the specificity of each phosphorylation changes in the AD and non-AD groups, we assessed linear correlations between each phosphorylated peptide and its unmodified version (Fig. 2). Strong correlations were found for all investigated phosphorylated sites in the AD group (R^2^ ranging from 0.76 to 0.94, Fig. 2) and non-AD group (R^2^ ranging from 0.66 to 0.91) except for pT217, showing broader diffusion of participant’s point values (Fig. 2-B) and pT205, which was not detected in almost all participants (Fig. 2-D). Interestingly, pT217/T217 ratio and slope comparisons demonstrated a highly significant hyper-phosphorylation rate in the AD population versus non-AD group (slope increases 2.92 times, p<0.0001, Fig. 2-B), providing the first evidence of an increase in its phosphorylation stoichiometry in AD patients. In contrast, a hypo-phosphorylation at S199 and S202 (slope decreases 2.66 times for S199 and 3.63 times for S199+S202, p<0.0001, Fig 2-C to 2-G) was observed. This relative hypo-phosphorylation was concomitant to the appearance or increase in T205 phosphorylation (slope increases 11 times, p<0.0001, Fig. 2-D, E, G). The pT181/T181 ratio and slopes of comparisons indicated only a slight increase in T181 phosphorylation in the AD versus non-AD groups (slope increases 1.46 times, *p*=0.02, Fig. 2-A) mainly driven by the values of AD patients with the highest CSF tau levels.

**Fig. 1.**
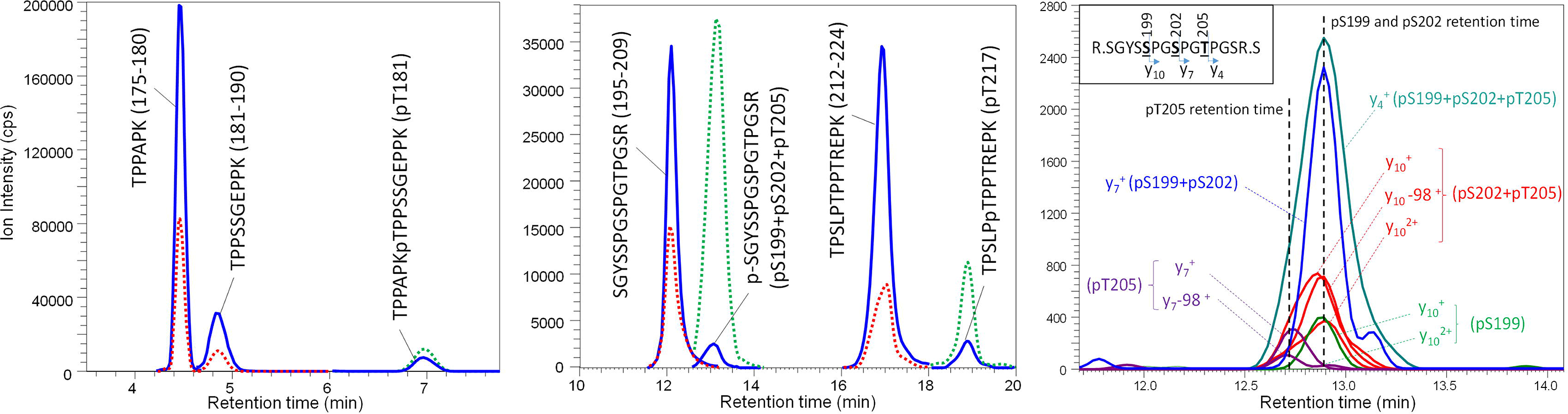
Quantitation of phosphorylated tau isoforms in CSF. Sum of Extracted Ion Chromatograms from Parallel Reaction Monitoring (PRM) analysis of tau phosphorylated peptides and the corresponding unmodified peptides in CSF. **Panel A**, T181 monitoring using a microLC system. **Panel B**, S199, S202, T205 (coeluted in framed signal) and T217 monitoring using a nanoLC system. Endogenous signals (full blue line), 15N labeled peptides (red dotted line), AQUA peptides (green dotted line). **Panel C** shows specific PRM transitions, according to Biemann nomenclature for peptide fragmentation, which allowed identification of the three coeluted monophosphorylated peptides carrying pS199, pS202 or pT205. Cps= Count per second.

**Fig. 2.**
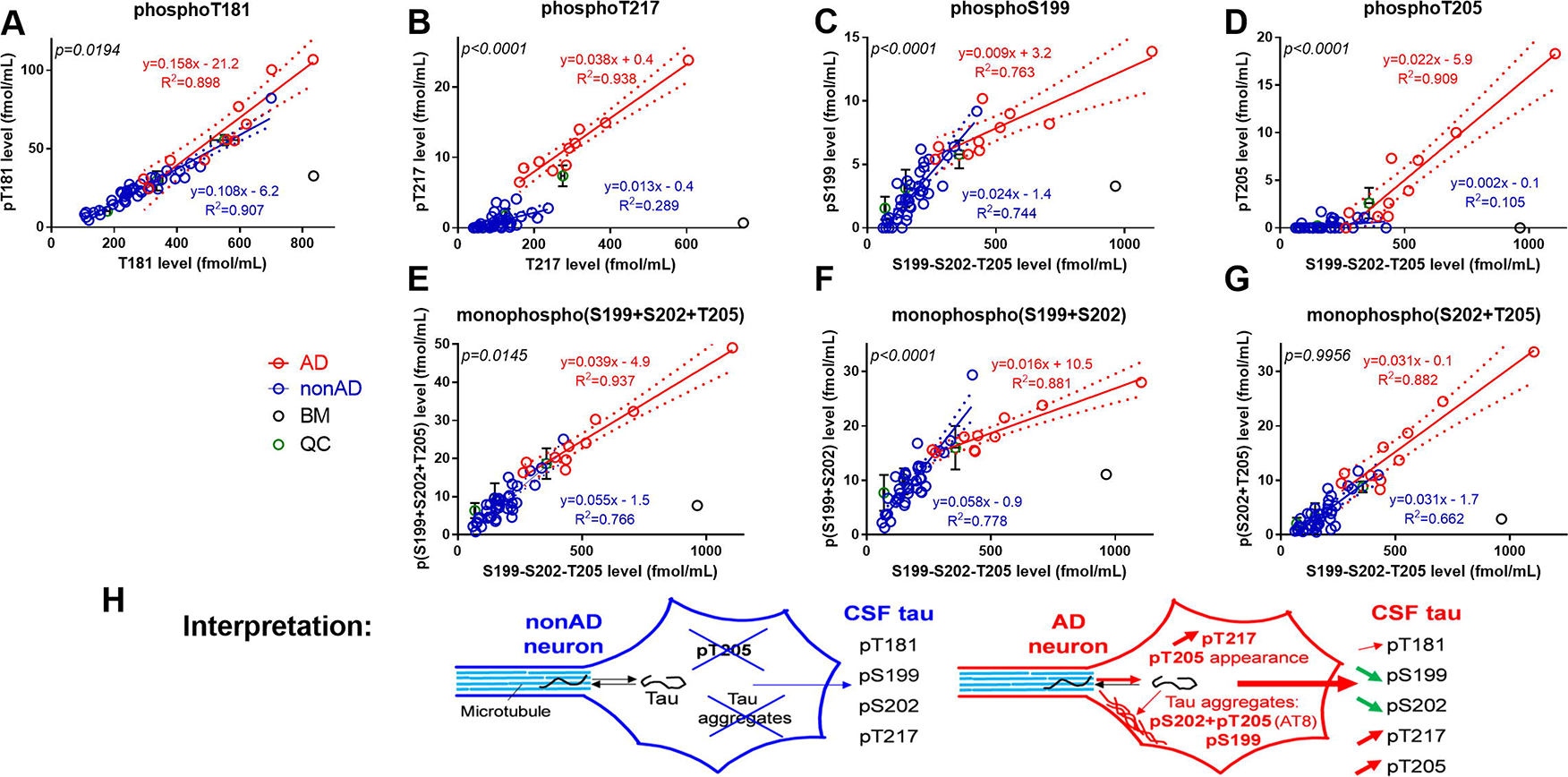
Hyper-phosphorylation and hypo-phosphorylation of CSF tau, assessed by comparing stoichiometric ratios on phosphorylated sites in discovery cohort. The slopes of phosphorylated isoforms to non-phosphorylated isoforms indicate the relative site phosphorylation extent in AD (red) and non-AD (blue). Relations between mono-phosphorylated peptide levels and their unmodified counterparts (**Panels A to D**). Least square regression models are plotted for AD and non-AD. 95% confidence intervals and p-values for slope difference comparisons are shown, indicating specific hyper-phosphorylation of pT217 and pT205, hypo-phosphorylation of pS199 and subtle hyper-phosphorylation of pT181. **Panels E-F-G** show sums of monophosphorylated peptides levels from the 195-209 tau sequence measured using shared transitions. Results and corresponding standard deviation from three quality controls (QC) are shown. **Panel H**: Schematic interpretation on differential levels of phosphorylated tau species in neurons and CSF from non-AD and AD patients.

### 3.2 Relevance to AD diagnosis of site-specific CSF tau phosphorylation

The specific T217 hyper-phosphorylation discriminated AD and other diseases or cognitively normal controls (Fig. 3-A). The sensitivity and specificity of pT217/T217 MS ratio (AUC 0.995) were higher than those of the ELISA p-tau(181)/t-tau (AUC 0.691) and MS pT181/T181 (AUC 0.856) ratios to discriminate AD versus non-AD (Fig. 3-B). Six subjects from the non-AD clinical group presented higher T217 phosphorylation rate (close to or similar to AD), compared with the other non-AD subjects (Fig. 3-C, ratio>1.5%, p<0.0001). Clinically, they were classified as LBD (n=3), VD (n=1) or AD with concomitant cerebrovascular diseases (AD with CVD) (n=1) and ACIH (n=1). They had normal CSF tau concentration, assessed by ELISA or MS, but a low Aβ42 level (Fig. 3-D), suggesting the presence of amyloidosis in the brain. The AD patient with CVD showed a high T217 phosphorylation rate supporting the presence of an AD underlying process. The comparison of the three LBD patients with enhanced T217 phosphorylation and the six others LBD patients revealed no demographic, clinical or biological profile differences.

**Fig. 3.**
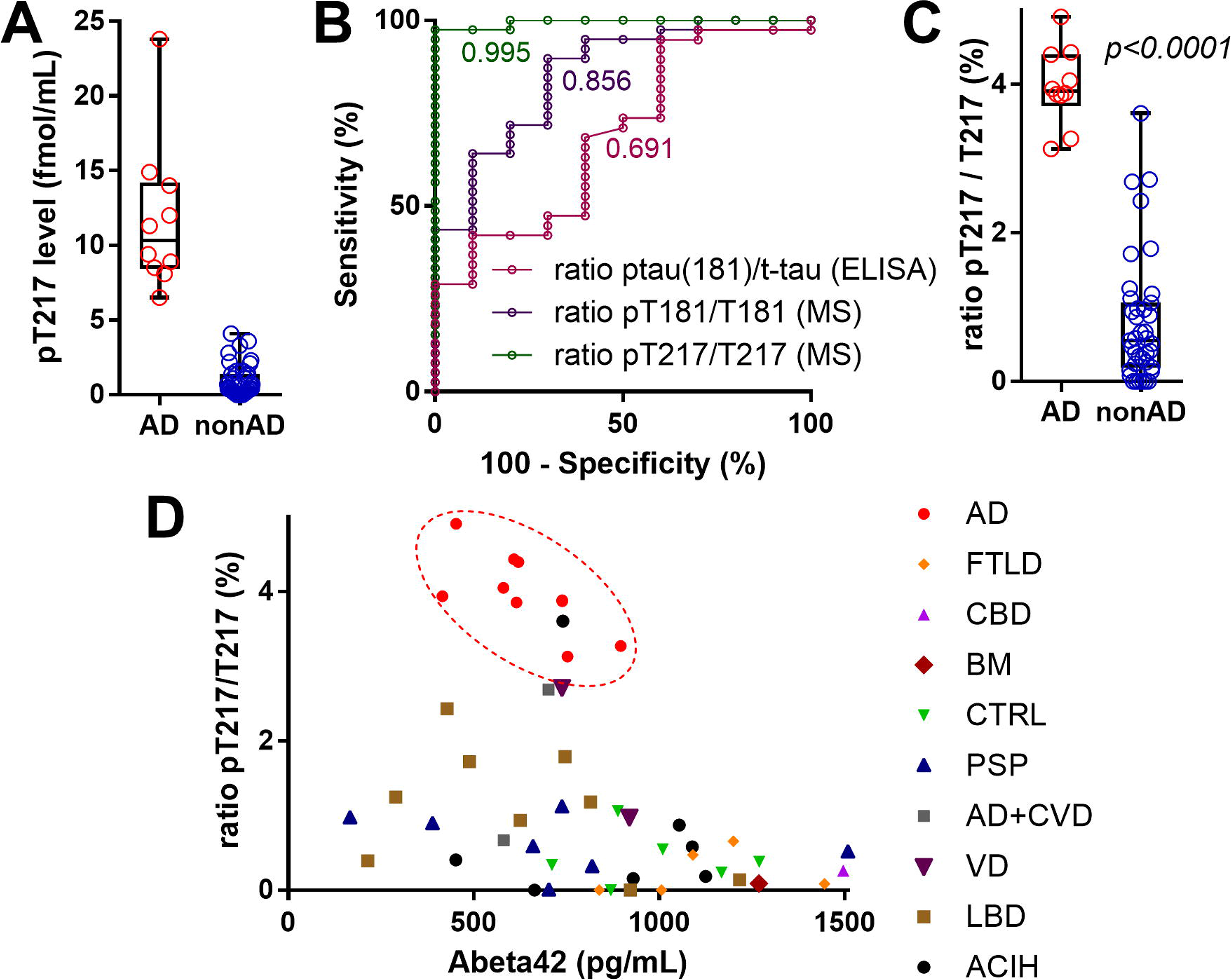
CSF tau phosphorylation on T217 is a specific diagnostic biomarker for AD. **Panel A**: T217 hyper phosphorylation discriminates AD participants (mild and moderate AD) from disease controls. Higher levels of pT217-containing peptide observed in AD result from the combination of T217 hyperphosphorylation and increase in CSF tau level. **Panel B**: ROC curves for the diagnosis of AD from non-AD participants using phosphorylation rate of T217, T181 by MS and T181 by ELISA. **Panel C**: Comparison of pT217/T217 ratio values obtained in AD and non-AD groups indicates a specific hyper-phosphorylation in AD on T217. **Panel D**: Distribution of pT217 rates and Aβ42 levels measured by ELISA across the participants. The AD group cluster has the highest pT217/T217 ratios and low Aβ42 (dashed red circle). Positive amyloidosis was considered for Aβ42 levels <750pg/ml.

### 3.3 Validation in a second cohort

We next validated the increased phosphorylation state of tau at T217 in AD in a second cohort from the Knight AD Research Center (ADRC) at Washington University in Saint Louis comprising 86 participants with no cognitive complaint or mild cognitive impairment (16). Consistent with the results of PiB-PET and CSF A 42/A 40 ratio measured by MS, participants were stratified in 29 amyloid-positive, 47 amyloid-negative participants and 10 cases with conflicting results between PET-PIB and CSF profile. To improve the assay sensitivity, immunopurification with the Tau1 antibody was performed (Sato et al, in preparation). Isotopically labeled versions of both phosphorylated and unphosphorylated peptides were used to measure site phosphorylation ratio. All phosphorylated peptides targeted were detected in all samples, with the exception of pS199 containing peptide not recovered by the Tau1 antibody. In this cohort, we confirmed the positive diagnosis relevance of pT217 biomarker at early stage of the disease. The pT217/T217 ratio clearly separated amyloid positive and negative groups (Fig. 4-B and 4-C, AUC 0.999). In addition, the measurement of pT181/T181 ratio discriminated the amyloid-positive and amyloid-negative groups (Fig. 4-B, AUC 0.956) consistent with the tendency observed in the discovery cohort (Fig. 3-B, AUC 0.856). T217 and T181 phosphorylation ratios were also correlated, (r=0.524, *p*=0.0002, SI Appendix, Fig. S5), suggesting that the phosphorylation of these sites could result from a common pathway. However, the diagnosis sensitivity of pT181 was lower than that of pT217. Both phosphorylation ratios were better discriminators than p-tau(181) and t-tau levels measured by ELISA (AUC 0.874 and 0.932 respectively, SI Appendix, Fig. S6).

**Fig. 4.**
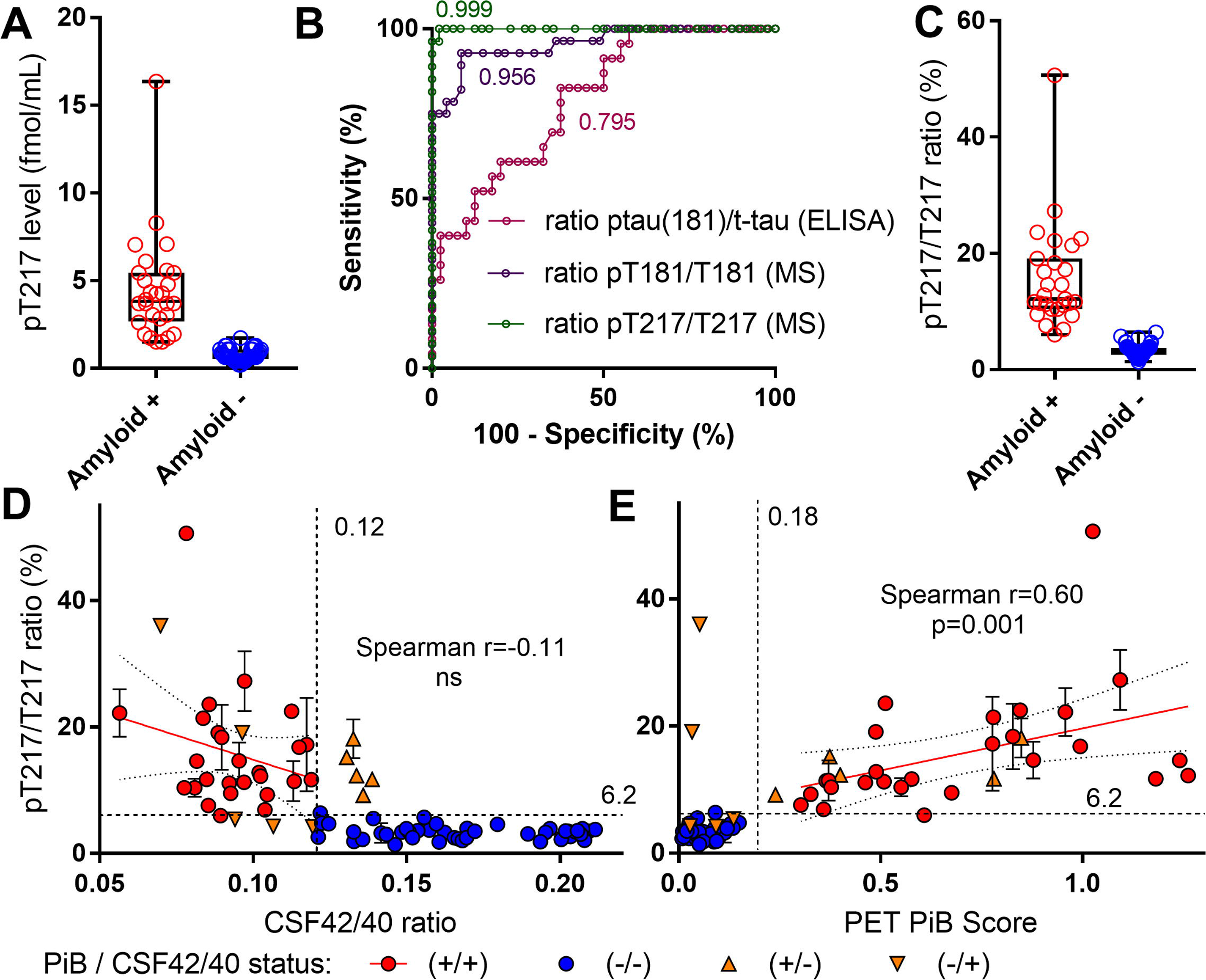
CSF tau phosphorylation on T217 is associated to amyloidosis status. **Panel A**: T217 phosphorylation significantly increases in participants having amyloidosis (PiB-PET and CSF42/40 ratio positive) compared to amyloid-negative controls with no or mild cognitive decline. **Panel B**: ROC curves for the diagnosis of amyloid-positive from amyloid-negative participants using phosphorylation rate of T217, T181 by MS and T181 by ELISA. **Panel C**: pT217/T217 ratio comparison demonstrates the specific phosphorylation on T217 in participants with amyloidosis. **Panel D-E**: Comparison of T217 phosphorylation with CSF Aβ42/40 changes measured by MS and to amyloid plaque deposition measured by PiB-PET. **Panel D**: The extent of T217 hyperphosphorylation is not correlated with the decrease of CSF Aβ42 relative to Aβ40. The five conflicting cases (orange triangles, positive for PiB-PET and T217 hyperphosphorylation but negative for CSF Aβ), were all slightly above the 0.12 threshold chosen to define the amyloid status, suggesting that they may result from insufficient sensitivity of the CSF amyloid assay. **Panel E**: PiB-PET loading (FBP Total Cortical Mean) is correlated with T217 phosphorylation state in amyloid-positive participants. Cut-off value differentiating amyloid positive from amyloid negative by PiB is 0.18.

### 3.4 Correlations between T217 hyperphosphorylation, amyloid pathology and cognitive status

We next determined the correlations between this new biomarker and the amyloid process. In the Montpellier cohort, we found a significant correlation between T217 hyperphosphorylation and CSF Aβ42 level (*p*=0.003, r=-0.42), suggesting a potential relationship between T217 hyperphosphorylation and the amyloid pathway in AD patients at mild to moderate stage of the disease. Our results in the ADRC cohort also suggested a strong relationship between T217 hyperphosphorylation and amyloid status. Importantly, T217 phosphorylation rate was significantly correlated with the extent of PiB-PET measured by FBP Total Cortical Mean (Fig. 4-E, r=0.60, *p*=0.001). Moreover, all the five conflicting cases with PiB-PET(+)/CSF(-) amyloidosis profiles were hyperphosphorylated on T217 (Fig. 4-D and 4-E). In contrast, no significant correlation was found between T217 phosphorylation and CSF Aβ42/Aβ40 ratio. Discrepancy between PiB-PET and CSF Aβ42/Aβ40 could be attributed to an insufficient sensitivity of the CSF amyloid assay, as corresponding CSF Aβ42/Aβ40 MS ratios were close to the threshold of quantification (0.12 ± 0.02, Fig. 4-E). Among the five participants with opposite PiB-PET(-)/CSF(+) amyloidosis, two exhibited hyperphosphorylated T217 (Fig. 4-D and 4-E). This suggests these cases highlighted by their high tau hyperphosphorylation could have significant CSF Aβ changes prior to amyloid plaques deposits detection. Combination of CSF Aβ42/Aβ40 ratio with T217 phosphorylation rate confidently identifies amyloid-positive participants without PiB-PET data even with CSF Aβ42/Aβ40 ratio values in the intermediate range slightly above the threshold (0.12 ± 20%). Amongst participants with no cognitive complaint (CDR-SB=0), T217 phosphorylation was able to completely distinguish amyloid-positive (n=9) from amyloid-negative participants (n=26, AUC 1.00, Fig. 5), further supporting the ability of this marker to confidently identify preclinical AD participants. However, no correlation was observed between global cognitive performances measured with CDR-SB and T217 phosphorylation ratio in this population (Fig. 5).

**Fig. 5.**
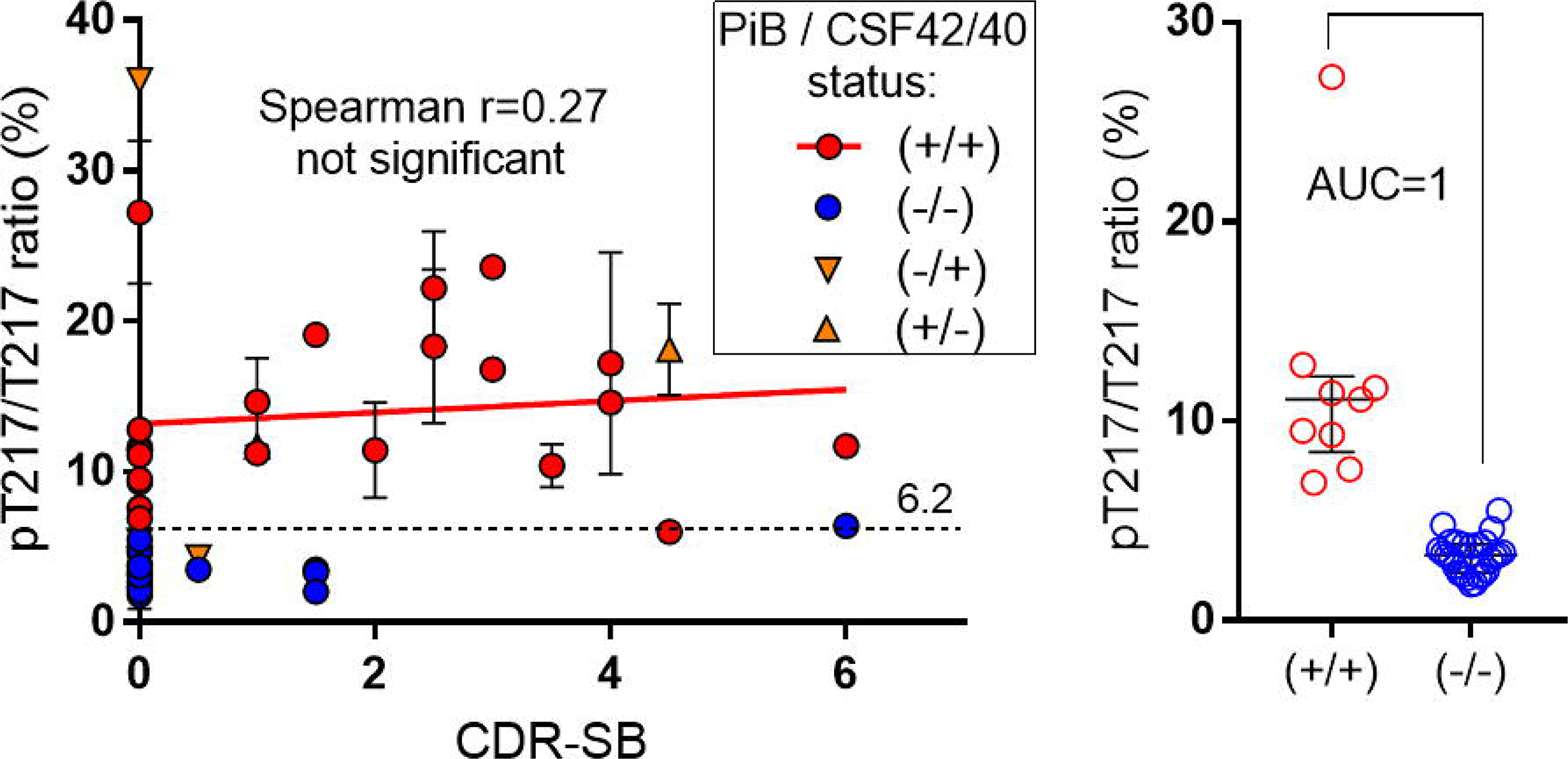
CSF tau phosphorylation on T217 is independent from cognitive status and is significantly modified in preclinical AD. Left Panel: No correlation exists between T217 phosphorylation and the cognitive profile measured by the clinical dementia rating sum of boxes (CDR-SB). Right Panel: Amongst participants with no cognitive decline (CDR-SB=0), T217 is already significantly hyper phosphorylated in the amyloid-positive group.

## 4. Discussion

Using innovative targeted-HRMS methods simultaneously measuring low-abundant phosphorylated tau peptides and their unmodified counterparts in CSF, we provide for the first-time direct evidence of changes in CSF tau phosphorylation rates concomitant with amyloid-β changes and distinct from increased CSF tau concentration. This approach allows for studying AD-specific tau phosphorylation rates assessing hyper- and hypo-phosphorylation to indicate underlying abnormal metabolism.

The most striking result of the current study is the highly specific hyperphosphorylation of T217 in CSF tau from preclinical, mild and moderate AD participants investigated in two independent and well-characterized cohorts. Amyloid-β pathology, which likely occurs before AD tauopathy, and tau hyperphosphorylation on T217 were strongly associated throughout the disease stages. This supports the hypothesis that amyloidosis can be related to tau phosphorylation changes (17). The absence of participants with β-amyloidosis and normal T217 phosphorylation in this study suggests that this tau phosphorylation site could follow the amyloid process. The hyperphosphorylation of T217 could be a key player in the AD pathophysiological process and its role differed from other tau biomarkers. In our cohorts, the increase in the levels of CSF tau or p-tau, as assessed by ELISA, was less effective in identifying amyloidosis status than pT217/T217 ratio. The specificity of these AD tau biomarkers may improve the prediction of the risk of cognitive decline in individuals who ultimately progress to clinical AD (18). Thus, combining amyloidosis biomarkers with pT217/T217 ratio measurement to detect AD, should dramatically improve diagnosis of preclinical AD. Although T217 hyperphosphorylation is highly correlated to amyloid plaques measured with PiB-PET load than amyloid markers in the CSF, tau modification may not be exclusively caused by the presence of fibrillary plaques. For the two participants with T217 hyperphosphorylation and low CSF Aβ 42/40 without brain amyloid load (Fig. 4-E), they could be cases with diffuse plaques or Aβ oligomeric formation only, not detected by PiB-PET (19) (20) but contributing to reduce CSF Aβ42 levels (21).

The wide-spread use of pT181 as an AD marker in clinics is likely due to its high level of detectability, but not necessarily high specificity (22). Measurement of pT181 highlights a slight change in its phosphorylation stoichiometry in AD, compared with non-AD dementia. The increased phosphorylation rate of T181 was even more apparent in a well-characterized amyloid-positive group investigated in the validation cohort. In both studies, the ratio combining ELISA t-tau and p-tau181 failed to demonstrate the change of phosphorylation ratio occurring on T181, stressing the limitations of ELISA to monitor accurately these changes. Thus, the increasing of pT181 broadly reported in AD CSF mainly results from the concomitant increasing of tau isoforms level rather than significant change on the T181 phosphorylation stoichiometry. Amyloid-β pathology induces hyperphosphorylation on T181 but the resulting modification appears less specific or significant in comparison to the relative increasing observed on T217.

Modifications of the phosphorylation stoichiometry observed on S199, S202 and T205 in AD CSF likely reflect specific changes in tau phosphorylation previously observed in AD brain. Indeed, pS202 and pT205 are part of the doubly phosphorylated epitope recognized by the AT8 antibody (33). AT8 binds to tau aggregates found in AD brain autopsies and the extent of AT8-immunoreactive aggregates correlates with the severity of tauopathy (Braak stages) (23-26). AT8 has no reactivity for normal tau or tau related to AD measured in CSF (27). Moreover, the Tau1 antibody that binds to normal tau at a non-phosphorylated epitope containing S199 has no affinity for AD tau aggregates (28). Together, these findings support a concomitant and abundant hyperphosphorylation of S199, S202 and T205 in tau aggregates from AD brain (29). However, controversy exists about the intrinsic property of such phosphorylated tau isoforms to promote tau aggregation in AD (30). The decreased amounts of S199 and S202 phosphorylation observed in mild and moderate AD CSF from the first cohort is consistent with an accumulation of the corresponding phosphorylated tau isoforms in aggregates (Fig. 2H). Furthermore, the detection of T205 phosphorylation mainly in moderate AD CSF suggests an important role of this site in pathological mechanisms underlying amyloid-β related tauopathy. Modification in S202/T205 phosphorylation was not detected in the second cohort composed of preclinical and mild AD participant (data not shown), suggesting a dynamic process of tau phosphorylation on specific sites during the course of the disease.

The present findings could suggest an interplay between AD amyloidosis and hyperphosphorylation of tau on T217 and, to a lesser extent, on T181. These sites are both substrates for the serine/threonine proline-directed kinase GSK-3β, and the activation of GSK-3β by Aβ oligomers has been proposed as a link between amyloid peptide and tau phosphorylation (31). The common and relatively well-correlated hyperphosphorylation on these two GSK-3β sites in patients with amyloidosis could be consistent with such a mechanism. Further studies designed to compare tau phosphorylation rates in CSF and brain, including tau aggregates measured by tau PET, are likely to provide novel insights into AD pathophysiology and may identify novel therapeutic approaches. These findings point to a specific link between amyloid plaques and tauopathy in AD and provide a potential link in the cascade of molecular events that leads to AD. Thus, given the specificity of pT217 within the AD process, it may represent an important target for future therapeutic development and an interesting tool to follow potential effects of anti-amyloid drugs in limiting this abnormal tau metabolism.

## ACKNOWLEDGMENTS

We thank Serge Urbach and Martial Seveno for their assistance in MS experiments performed at the FPP of Proteomics Pole of Montpellier-Languedoc-Roussillon. Vitaliy Ovod for his work in assembling clinical data for the validation study and Paul Moiseyev for his assistance. Nicholas Kanaan for Tau1 antibody. Isabelle Huvent for 15N Tau recombinant protein. Joel Ménard for his continuing support in the ProMarA study. Yves Dauvilliers for advices regarding redaction of the manuscript. This work was supported by the 2010 National PHRC “ProMarA” (S.L.), the Alzheimer’s Association Research Fellowship (N.R.B), by the NIH R01 NS065667 and NS095773 grants (R.J.B.) and the Coins for Alzheimer’s Research Trust Fund (C.S.).

